# Profiling the size-dependent heterogeneity of membrane proteins in a mixed population of small extracellular vesicle for potential cancer diagnosis

**DOI:** 10.1101/2022.04.13.488159

**Authors:** Chunhui Zhai, Feng Xie, Qiang Zeng, Weiqiang Zheng, Jingan Wang, Haiyan Hu, Yuting Yang, Xianting Ding, Hui Yu

## Abstract

The heterogeneity in small extracellular vesicles (small EVs) introduces an extra level of complexity in small EV-based liquid biopsy for cancer diagnosis. Heterogeneous membrane protein expression is correlated with sizes of small EVs, but accessing this correlative information is limited by the precise isolation of size-dependent subpopulations. Herein, we present a single EV enumeration (SEVEN) approach to profile protein heterogeneity in size-dependent subpopulations, and demonstrate its potential in improving the accuracy of cancer diagnosis. The interferometric plasmonic microscopy (iPM) capable of imaging single biological nanoparticles with the diameter down to 30 nm is employed to detect small EVs at the single-particle level. Small EVs population with mixed sizes are directly imaged, individually sized and digitally counted during their binding onto different aptamer-coated iPM sensor surfaces. The protein expression levels and binding kinetics of three size-dependent subpopulations are analyzed, forming a multidimensional data matrix for cancer diagnosis. Using small EVs derived from different cancer cell lines, highly heterogeneous protein profiles are recorded in the three subpopulations. We further demonstrate that the cancer classification accuracy could be greatly improved by including the subpopulation level heterogeneous protein profiles as compared with conventional ensemble measurement.

## Introduction

Extracellular vesicles (EV) are a group of membrane-enclosed phospholipid vesicles secreted by mammalian cells, including normal cells as well as cancer cells^1^. Small EV (sEV), including exosomes and a portion of microvesicles, is a unique subset with a diameter of less than 200 nm^2^. EVs, including sEV, are packaged with functional molecules (i.e. proteins, amino acids and nucleic acids) of their parental cells, and play an important role in cell-cell communication^3^, immune response^4^ and cancer metastasis^5 6^. sEV are highly heterogeneous in the membrane proteins, sizes and contents depending on the cell sources, cancer-gene mutation and other environmental factors^7 8 9^, making them a potential tool in cancer diagnosis^10 11^. For example, by profiling the expression level of membrane proteins from sEV, early diagnosis of breast cancer^12 13^ and classification of cancer types^14^ have been demonstrated.

With sophisticated isolation techniques, size-dependent subsets of sEV including exomere (<50 nm), Exo-S (60 - 80 nm) and Exo-L (90 - 120 nm) were identified with unique molecular biomarker profiles^15^. This suggests that conventional analytical approaches by measuring average information from the full population of sEV would inevitably suffer from the large background noise from irrelevant subpopulations^16^. Profiling protein heterogeneity of the size-dependent sEV subpopulations would thus largely advance our knowledge in the mechanism of their biological functions, as well as the development of accurate diagnostic tools^8^. However, the major challenge to access the heterogeneity information is the difficulties in precisely isolation of sEV subpopulations with specific sizes. Although there are several emerging techniques for isolation of sEV subpopulations ^15 17 18 19 20 21 22^, they typically suffer from poor efficiency, time consuming process, and sophisticated fabrication^*23*^.

Herein, we present an alternative approach termed single EV enumeration (SEVEN) to profiling the protein heterogeneity in size-dependent subsets of small EVs for cancer diagnosis. Instead of profiling proteins on isolated size-dependent subsets in the conventional approaches, SEVEN accurately sizes single small EVs captured by different aptamers to access the heterogeneity information. Using small EVs derived from five different cancer cell lines, we measured the heterogeneous expression of five different membrane proteins on three small EVs’ subsets, and developed a machine learning algorithm for cancer classification.

## Results and Discussions

### The overview of SEVEN

The principle of SEVEN is based on our previous work to image, size and digitally count single sEV by the interferometric plasmonic microscopy (iPM)^24 25^ (**Figure 1**). For each protein biomarker detection, SEVEN dynamically measures the size of each sEV particle specifically binding onto an aptamer-coated sensor surface (**Table S1**), and divides them into size-dependent subpopulations. Corresponding size-dependent binding curves are then constructed by digital counting of sEV, from which parameters including the maximum binding number and the exponential coefficient are quantified. For sEV from different cell sources measured on different aptamer-coated sensor surfaces, these parameters form a multidimensional matrix containing size-dependent heterogeneous information of protein biomarkers.

**Figure 1.**
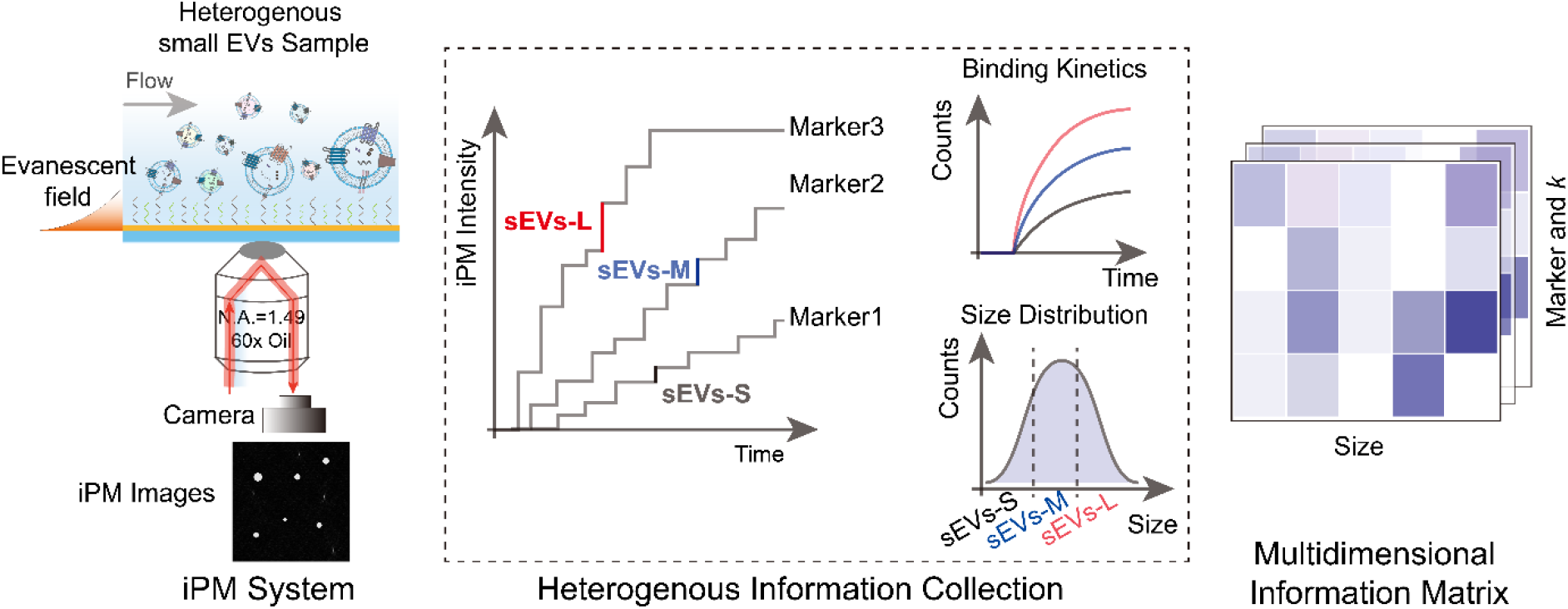
The schematics. Binding of sEV onto specific aptamer-coated sensors is imaged with the iPM system; image intensity and counts are analyzed to construct binding curves of each subpopulation (sEV-L, sEV-M and sEV-S); a matrix is formed with the sizes, counts and kinetics.

### Verification of sEV sample

Five different cell lines, including A549 (lung cancer), HepG2 (liver cancer), MCF-7 (breast cancer), LNCaP (prostatic cancer) and L-02 (normal liver cells), were cultured to derive sEV. sEV were isolated from the culture medium using ultracentrifugation methods as described previously^26^. The sizes of isolated EVs were below 200 nm as measured by nanoparticle tracking analysis (NTA) (**Figure 2a** and **Figure S1**), and the concentration varied from 10^10^ /mL to 10^12^ /mL. The morphology of sEV was characterized by transmission electron microscopy (TEM), showing typical entire and saucer-like shapes (**Figure 2b**). According to the guidelines in minimal information for studies of extracellular vesicles 2018 (MISEV 2018)^27^, we found that three universal EV-positive plasma membrane proteins, Alix, CD63 and Tsg101 were positively expressed, and one EV-negative protein, Calnexin, was negatively expressed with western blot (**Figure 2c**).

**Figure 2.**
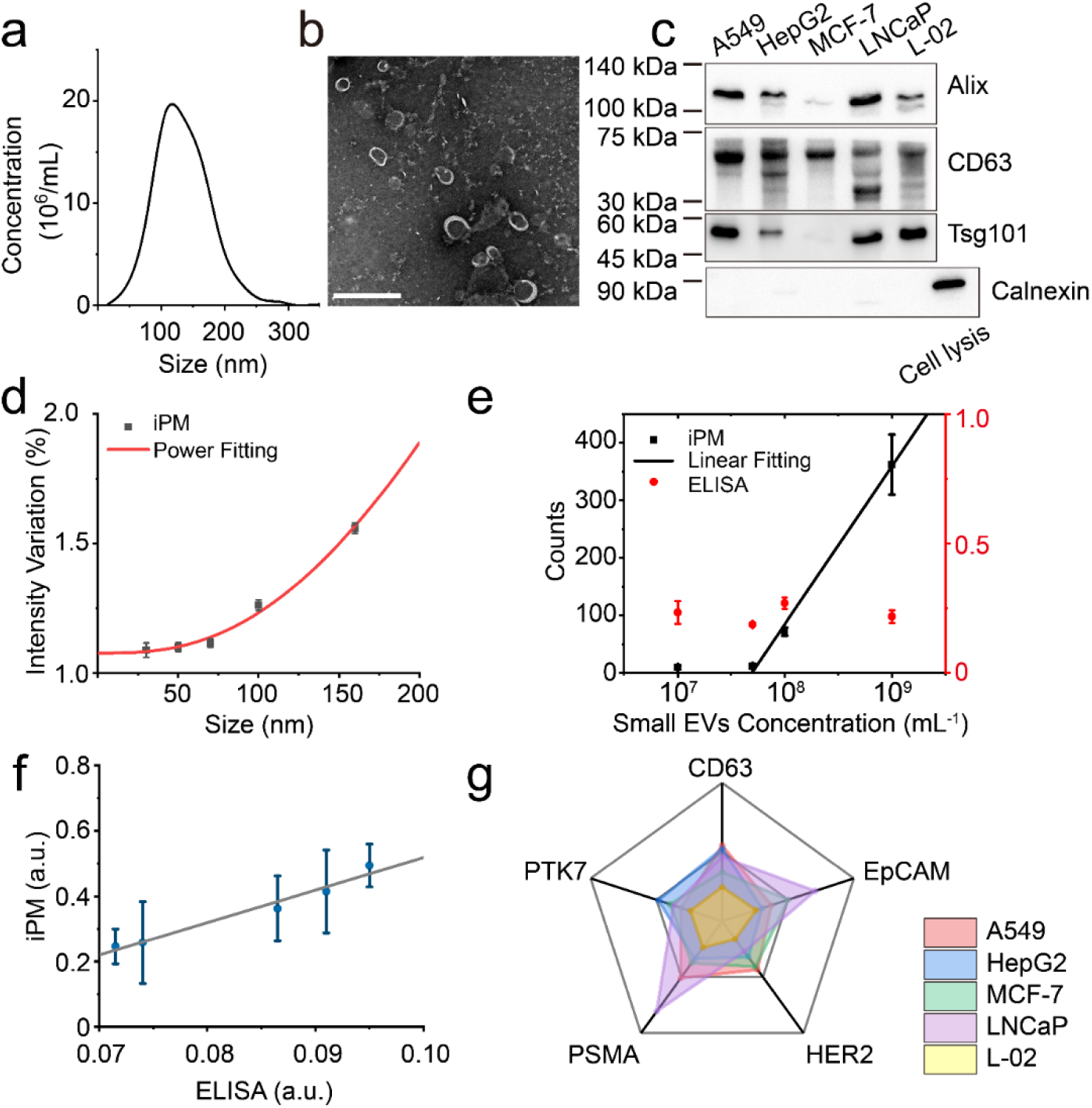
Single sEV imaging and detection. (a) Concentration and size distribution of A549-derived sEV measured by NTA. (b) Typical TEM images of sEV. (c) Western blot results of Alix, CD63, Tsg101 and Calnexin in sEV derived from A549, HepG2, MCF-7, LNCaP, L-02 cell lines. (d) Theoretical (red) and experimental (black) iPM intensities of silica nanoparticles of 30 nm, 50 nm, 70 nm, 100 nm, and 160 nm. (e) Sensitivity of iPM (black) and ELISA (red) for the detection of sEV from MCF-7 cells with CD63 aptamer -coated sensor chips. (f) CD-63 level on sEV of five cells lines determined by iPM and ELISA. (g) Radar plot showing iPM analyses of 5 EV protein markers from the five different cell lines (n = 3).

### Plasmonic imaging and detection of sEV

In SEVEN, the iPM system offers a unique approach to image and characterize single sEV. After binding onto the chips, sEV showed an intact morphology as characterized by scanning electron microscope (SEM) (**Figure S2**). We established the calibration curve between iPM intensity and particle sizes using silica nanoparticles (**Figure 2d** and **Figure S3**), and measured the sizes of sEV by intercalation. Using sEV samples from MCF-7 as an example, the number of EVs binding onto optimized HER2-aptamer modified sensor surfaces (**Figure S4**) within 15 minutes correlated well with the concentrations of total sEV in the range from 5 × 10^7^ /mL to 10^9^ /mL (**Figure 2e**). The specificity in measuring HER2-positive sEV was verified by comparing the binding on bull serum albumin (BSA)-coated surface and a dissociation experiment (**Figure S5**). Note that the conventional enzyme-linked immunosorbent assay (ELISA) failed to measure the HER2-positive sEV due to insufficient sensitivity.

Besides measuring the size and concentration, SEVEN could also report the average level of specific proteins in sEV samples. For sEV samples of the five cell lines at the concentration of 2 × 10^10^ /mL, we compared the CD63 expression levels measured by ELISA with the percentage of CD63-positive EVs in total EVs determined by iPM, which showed a good linear correlation (R^2^ > 0.96, **Figure 2f**). Similarly, the expression levels of protein biomarkers in the sEV from five cell lines were measured as the percentage of target-specific EVs, which were largely different from each other (**Figure 2g**). We note that the protein levels reported by SEVEN are not the expression level on single sEV, by related to the total proteins from the all sEV.

### Limitations in cancer diagnosis by total sEV analysis

We then explored the capability of SEVEN in profiling proteins in total sEV population for cancer diagnosis, which is the common practice. 72 small EV samples were collected from the cancer cell lines of A549, HepG2, MCF-7 and LNCaP, and 18 samples were collected from L-02 healthy liver cell lines as the control. The expression of the five biomarkers was first confirmed by Western blot (**Figure 2c** and **Figure S6**). Expression levels of CD63, EpCAM, HER2, PSMA and PTK7 in the small EV samples from the five cell lines were measured as the percentage of target-positive EVs in total EVs as described above (**Figure 3a**). The heterogeneity among different batches of small EV samples was obvious, which is inevitable due to the intrinsic variations in cell conditions and other environmental factors. The exponential coefficients related to the association rates were quantified, which were also heterogeneous among difference cell lines (**Figure 3b**).

**Figure 3.**
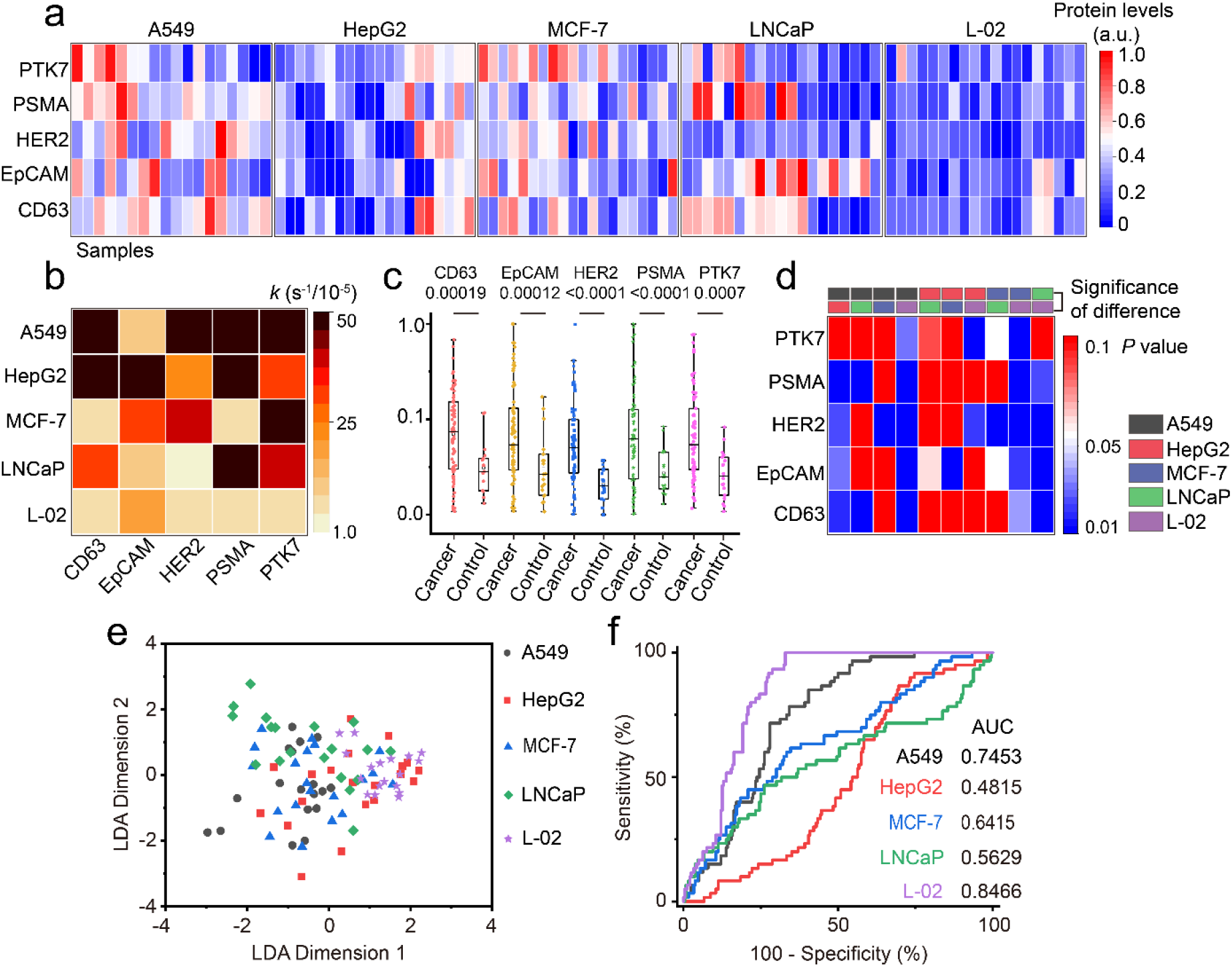
Profiling total sEV surface protein markers. (a) Heat map of sEV surface protein in 5 cell lines (18 samples for each cancer cell line). (b) The association rates of total sEV from five cell lines. (c) Elevated protein levels of all 5 protein markers in sEV derived from cancer cell lines (n = 78; means ± s.e.m.) than normal cell line (n = 18; means ± s.e.m.). (d) The statistical differences in distinguishing cancer types (n = 18 samples for each cell type) by a single biomarker by independent samples t-test. (e) Cancer classification using five protein biomarkers of total sEV by LDA. (f) The ROC curves for cancer classification in (e).

sEV from the four cancer cell lines showed a significantly higher protein levels than the healthy control from L-02 (*p* < 0.01, **Figure 3c**), indicating the potential to perform cancer diagnosis with these biomarkers. But when performing the cancer classification with the expression level and exponential coefficient of only one of the biomarkers, none was able to classify all five cancers with *p* < 0.05 (**Figure 3d**) in a pairwise comparison. Even the well-recognized specific markers, such as HER2 and PSMA, were not able to distinguish breast cancer and prostate cancer. When the expression levels and exponential coefficients of all five biomarkers were analyzed with the Linear Discriminant Analysis (LDA) algorithm used in previous study^13^, the overall average classification accuracy was only 34% (**Figure 3e** and **Figure S10 c**). The areas under the curve (AUC) were 0.74, 0.48, 0.64, 0.56 and 0.85 respectively, using receiver operating characteristic (ROC) analysis (**Figure 3f**).

These results suggest that using protein biomarkers in the mix population of sEV could lead to poor accuracy in the cancer diagnosis. One of the reasons could be due to the heterogeneity at the single EV level. For example, when examining the CD63 level in sEV from LNCaP cells using immunoelectron microscopy, different number of immuno-gold nanoparticles were observed on sEV (**Figure S7**). We then investigated the possibility to improve cancer diagnosis accuracy by exploring the protein heterogeneity at the subpopulation level.

### Profiling protein heterogeneity in size-dependent subpopulations

The sEVs were empirically divided into three size-dependent subpopulations, including the sEV-S (30-70 nm), sEV-M (70-120 nm) and sEV-L (120-160 nm). With the single EVs imaging and sizing capability of iPM, the binding curves of three subpopulations were digitally plotted (**Figure 4a** and **Figure S8**). The exponential coefficients were obviously heterogeneous (**Figure 4b**). For example, the exponential coefficients were 0.0070, 0.0091, and 0.0089 s^-1^ for sEV-S, sEV-M and sEV-L binding with PSMA-aptamer, and 0.0013, 0.0015, and 0.0028 s^-1^ with CD63-aptamer respectively, and LNCaP and L-02 derived sEV showed much faster binding rates than A549-derived sEV.

**Figure 4.**
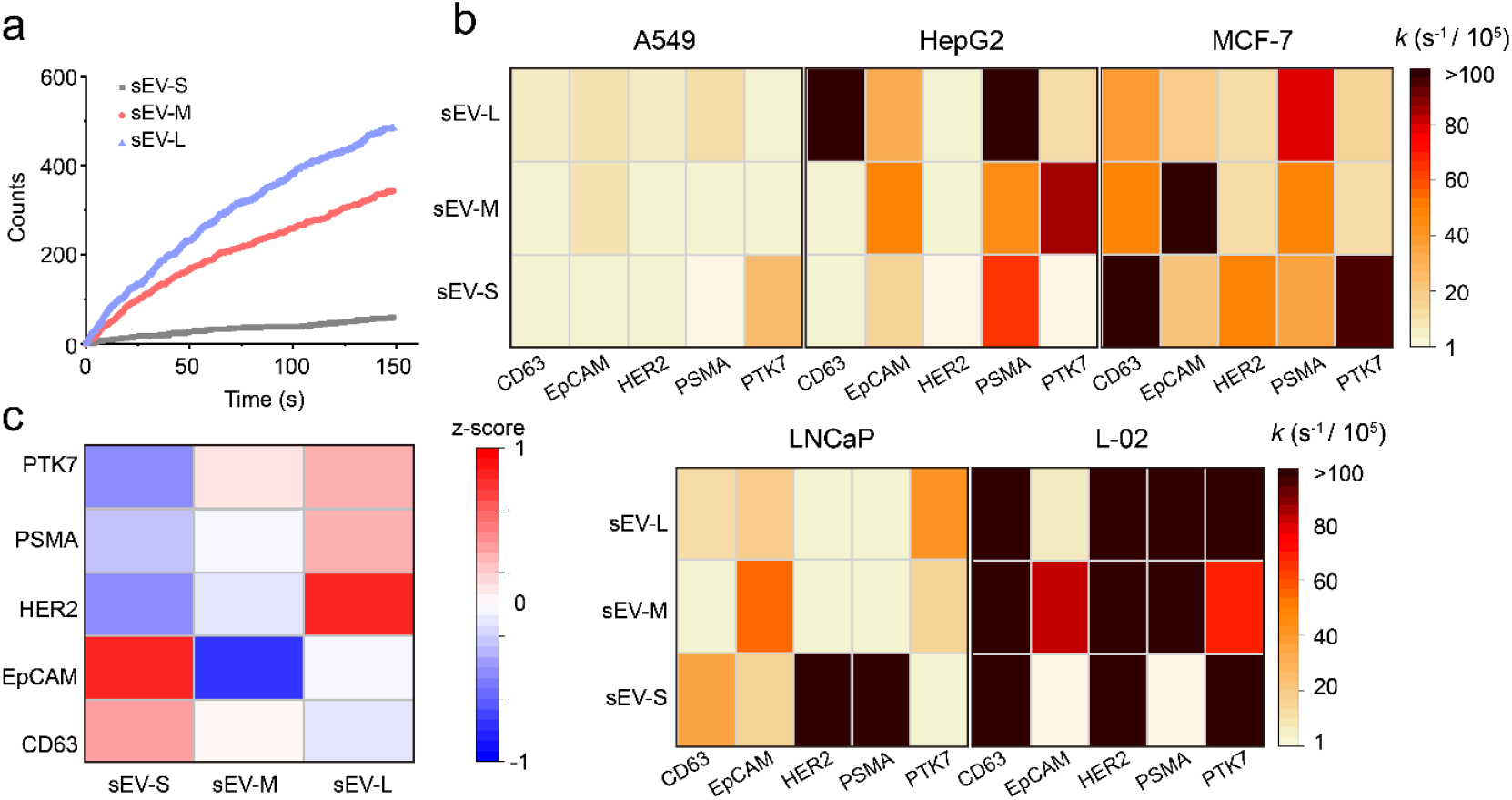
Binding kinetics and distinct marker of size-dependent sEV’s subpopulations. (a), An example of binding curves of LNCaP-derived sEV subpopulations on PSMA-modified chips. (b), The heat map of apparent association rate k of five kind of cell lines in five kind of aptamers-modified sensor chips. (c), Heatmap illustration of the relative abundance of sEV (A549) markers in sEV-S, sEV-M and sEV-L. Scale shown is protein levels subtracted by mean and divided by row standard deviation (that is, Δ (protein levels − mean)/s.d.).

We further calculate the z score of protein levels from the three subpopulations to highlight the difference^15^. In sEV from all cell lines other than A549, all protein markers were found to enrich in sEV-L (Figure S9). While for A549, EpCAM and CD63 were mainly expressed in the sEV-S subpopulations, but HER2, PSMA and PTK7 in sEV-L (Figure 4c). This confirms that membrane protein expression levels in sEV are not directly proportional to membrane areas, but instead, they were packaged purposely. The reason why specifically only EpCAM and CD63 from A549 cells were highly expressed in sEV-S is still unknown. Note that the sEV-s population weighed only a small portion of the total sEV (Figure 2a). Thus, when analyzing EpCAM and CD63 from the total sEV, the majority as sEV-M and sEV-L would give a large background noise, which explains the poor accuracy in Figure 3d.

### Improving cancer classification accuracy with subset information

The multidimensional information matrixes of sEV, including the protein levels and exponential coefficients of total sEV and those of three subpopulations were input into a LDA model to discriminate the five cell lines (**Figure 5a**). Using the same raw data and similar LDA algorithms in **Figure 3e**, the overall classification accuracy significantly improved to 70% (**Figure 5b, 5c**), with the AUC of 0.97, 0.85, 0.85, 0.97 and 0.82 for the five cell lines, respectively (**Figure 5d**). Precision Recall Curves (PRC) also showed a better performance of classification with subset information comparing to that without subset information to differentiate cancer cell lines (**Figure S10a, b**). Classification patterns of the HepG2 liver cancer cell line and the L-02 healthy liver cell line largely overlapped, but discrimination between cancer cell lines from different origins was reliable. When we combine the data of HepG2 and L-02 into a same group, the average accuracy was improved to over 81% (**Figure S10d**).

**Figure 5.**
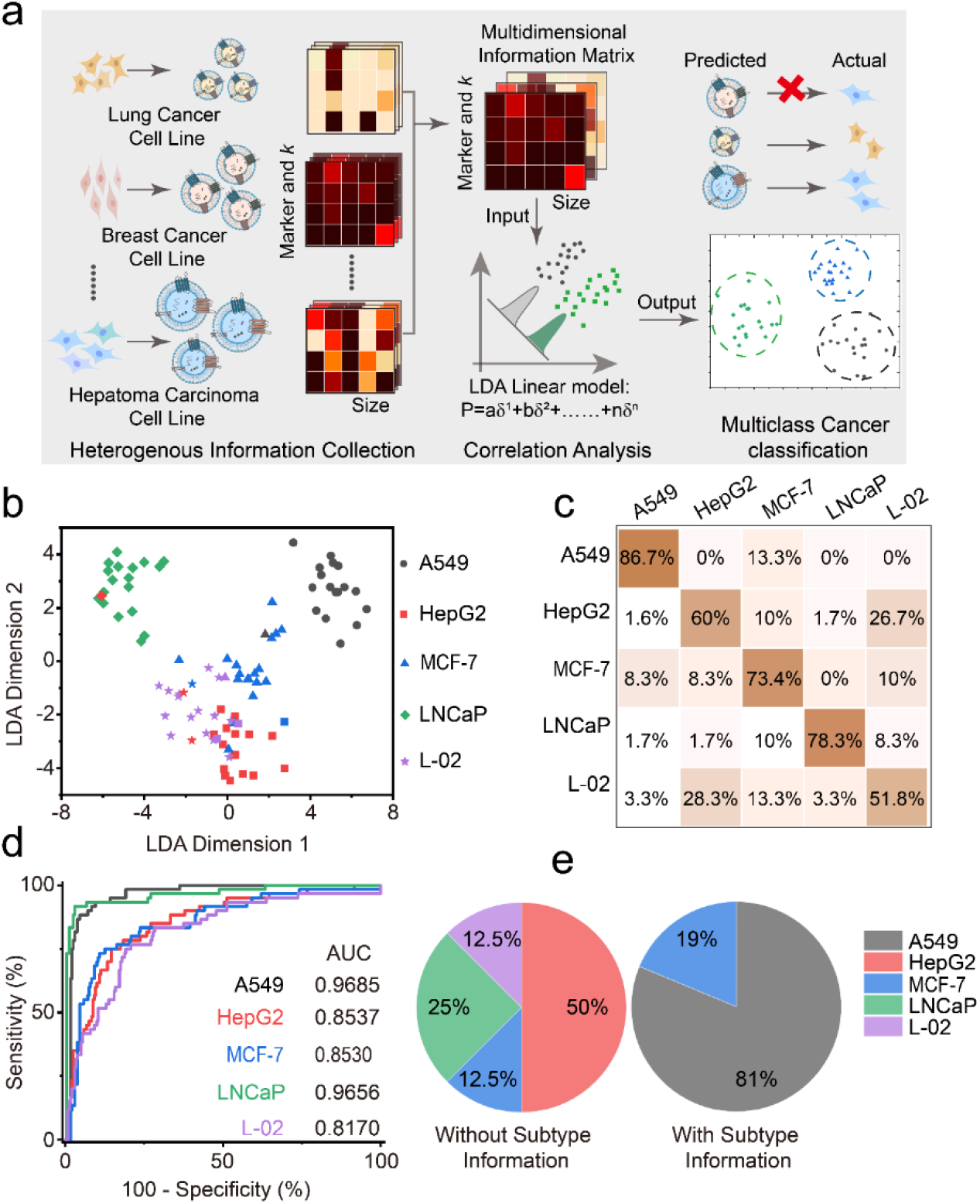
Multiclass cancer classification with subset information. (a) The workflow of SEVEN-based cancer classification with subset information. (b) The classification results by including subset information into Fig.3e. (c) Probability matrix summarizing the cancer classification results in (b). (d) ROC curves of (c). (e) Results in determining the origin of A549 derived sEV spiked in human serum samples.

We further performed simple validation experiments by spiking sEV derived from A549 cell lines into EV-depleted serum samples, and determined their origin with SEVEN. The ratio of success reached 81% (**Figure 5e**) with the information at the size-dependent subpopulations, while without this information none was correctly classified. This indicates the potential of SEVEN in analyzing sEV in complicated samples for future clinical applications.

### Re-defining the size-dependent subsets of sEV

Simply dividing the sEV into as small, medium and large subsets is intuitive but brutal, both in previous work by isolating the subsets and in this work by empirically setting thresholds. There has been little evidence showing why the membrane proteins and contents should be packed in such a simple size-dependent manner. A deeper study is hindered by the capability to isolate the sEV subsets within narrower size ranges. However, in SEVEN, the iPM system offers a sizing accuracy of 10 nm, which provides the opportunity to potentially address this problem. We thus re-calculated the SEVEN data between 30 to 160 nm as thirteen subgroups at the interval of 10 nm. Instead of empirically combining some of the subgroups, we developed an artificial intelligent algorithm to automatically search for the optimal size partition to achieve the best classification accuracy (Figure S11). The results show that when the sEV were grouped by sizes within 30-40 nm, 40-100 nm, 100-110 nm, 110-130 nm, 130-140 nm and 140-160 nm, the classification accuracy with raw data in Figure 5 dramatically increased to 87% (**Figure 6**). Besides, the healthy liver cell line L-02 was fully separated from cancerous cell lines.

**Figure 6:**
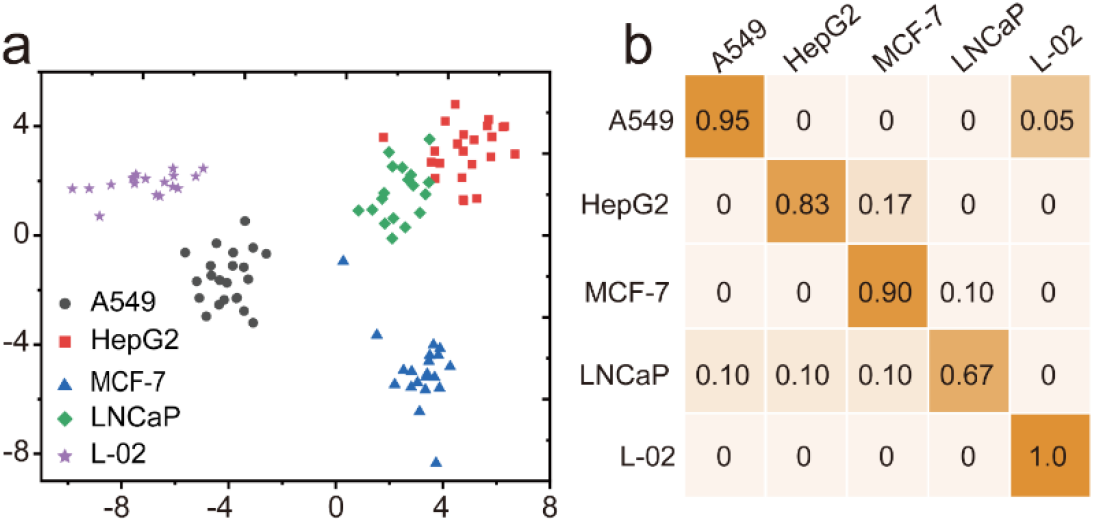
Multiclass cancer classification by re-grouping the subsets. (a) The classification results of five cell lines with raw data in Figure 5. (b) Probability matrix summarizing the cancer classification results in (a).

## Discussion

We have presented the single EV enumerating (SEVEN) approach to explore the in-depth information of size-dependent heterogeneity of sEV. Benefits from the single particle imaging and detection capability, SEVEN circumvents the challenges in precise isolation of subpopulations. Studies on five cell lines and five protein biomarkers have provided new evidence in the correlation between the protein levels and the size distribution of sEV. Experiments with sEV from cancer cell lines and those spiked in serum samples show that SEVEN could effectively improve the diagnostic accuracy over ensemble measurements. This work thus has highlighted the importance to explore the underlying relationship between different dimensions of heterogeneity of sEV for developing better diagnostic performance. On the meantime, there are still several limitations in the present work. First, the correlation between size and membrane proteins was obtained in a protein-by-protein manner, which is not ideal for this application. This could be improved by combining the total internal reflection fluorescence mode^28, 29^ and the iPM mode to simultaneously detect multiple membrane proteins and sizes at the single sEV level. Second, this work presented the first proof-of-concept study with cell lines and spiked samples. Further validation with clinical samples will be required to evaluate its potential in cancer diagnosis, which is currently undergoing.

## Materials and Methods

### iPM system

The iPM system was built on an inverted total internal reflection fluorescence microscope (TIRFM) (Olympus IX83) using a 60 × oil immersion objective (N.A. = 1.49). The surface plasmons were stimulated via Kreschmenn configuration by a laser beam at 637 nm (OBIS 637 nm LX FP 100 mW) at a highly inclined incident angle close to SPR dip angle. The real-time images of sEV were recorded by a sCMOS camera (Prime TM; Photometrics). A motorized XY stage (Ludl Electronic Products, Ltd.) was incorporated on the microscope to translate the sensor chip.

### iPM sensor chips and surface modifications

The sensor chips were 12-542-B (Thermo Fisher) glass coverslips coated with 3 nm of chromium and 47 nm of Au. The chips were cleaned first with deionized water and ethanol for three times, and dried by nitrogen gas. After a quick hydrogen flame treatment, the sensor chip was immediately immersed in modification buffer for 12 h. The modification buffer contained 1μM aptamer (Sangon Biotech, China) 5uM Tris (2-carboxyethyl) phosphine (TCEP, Sigma) and 1μM 6-Mercapto-1-Hexanol (MCH, Sigma). The chip was rinsed for three times with 1× PBS buffer to remove unbound aptamers, and 50 μL of bull serum albumin (BSA, Sigma) solution (1% w/v) was added to further block the residual active sites for 5 min. For positive-charge modification, the chip was treated with hydrogen flame and immediately submerged in 10 mM HS-PEG-NH_2_ (10,000 Da; Nanocs) water/ethanol (1:1) solution overnight.

### Image processing

The iPM images were processed offline with MATLAB R2018a (MathWorks). Raw images were preprocessed by moving average with n = 10 frames. The differential images between two adjacent average images were reconstructed with home-developed codes. Briefly, by calculating the radius and center of the ring in *k* space, the wave vector of single sEV was determined. Deconvolution was done in *k* space using the point-spread function obtained experimentally, by aligning 30 individual images of 100-nm silica nanoparticles to the maximum intensity point, and averaging after alignment. Then average intensity of the 3 × 3 pixels around the brightest pixel of particle images was calculated as the particle intensity.

### Size calibration

Silica nanoparticles (MikroNano Partikel GmbH) with the size of 30 nm, 50 nm, 70 nm, 100 nm and 160 nm were used to build the size-calibration curve. Raw silica nanoparticles were diluted with 1× PBS at 1:1000 vol/vol, and ultrasonicated for 30 min to redisperse the single particles followed by centrifuging at 2000 rpm to remove aggregates. 20 uL of nanoparticle solution was injected onto the positive-charge modified sensor chips, and images were recorded for 2 min at 100 fps. Each experiment was repeated in triplicates. The statistical histograms of silica nanoparticles were fitted with a Gaussian function to determine the peak intensity values (mean ± SD, n > 150) (Supplementary Fig. 4). The calibration curve was plotted as the intensity versus the diameter of silica nanoparticles (Fig. 2f).

### Cell culture

The human lung cancer cell line (A549, ATCC), hepatoma cell line (HepG2, ATCC), breast cancer cell line (MCF-7, ATCC), and normal liver cell line (L-02, ATCC) were cultured using high-glucose Dulbecco’s modified Eagle’s medium (DMEM) (Hyclone) with 10% extracellular-vesicle-free fetal bovine serum (EV-free FBS) (SeraPro) and 1% penicillin - streptomycin (Gibco). Prostate cancer cell line (LNCaP, ATCC) was cultured in Roswell Park Memorial Institute 1640 (PRMI-1640) (Hyclone) with 10% EV-free FBS (SeraPro) and 1% penicillin-streptomycin (Gibco). Cells were cultured at 37 °C with 5% CO_2_ in a humidified incubator (Thermo Fisher Scientific). The cells were incubated in the T75 flask (Corning) with 30% confluent and the culture medium was collected after 48 h culture when the cells were 70%-80% confluent.

### Isolation of small extracellular vesicles

sEV were isolated based on differential centrifugation. Cell culture media (300 mL) was first centrifuged at 500*g* for 5 min, followed by centrifugation at 2,000*g* for 45 min to remove cells. The treated medium was centrifuged at 10,000*g* for 60 min. Then the supernatant was filtrated by 0.22 μm membrane filtration (Millipore). Finally, the filtrate was ultracentrifuged at 100,000*g* for 120 min. The sEV were washed by 50 mL 1× phosphate buffer saline (PBS, Hyclone), followed by ultracentrifugation at 100,000*g* for 120 min. The purified sEV were resuspended in 50 uL 1× PBS.

### Spiking test in human serum

Human serum sample was collected from three healthy volunteers with agreements at the Shanghai Sixth People’s Hospital. Total volumes of 1 mL of clinical serum samples were first centrifuged at 2000*g*, and filtered through 0.22 μm pore filter (Millipore), then ultrafiltrated by 100 kDa ultrafiltration membrane (Millipore) at 10,000*g* at 4 °C for 20 min. The EV-depleted serum was stored at -80 °C before use. 10 uL A549 derived sEV was added to 90 uL EV-depleted serum to mimic the real cancer-related serum with the final EV’s concentration of 10^10^ / mL.

### NTA analysis

The size distribution of sEV samples were characterized by NTA (Particle tracking analyzer; Particle Metrix, PMX). All samples were diluted using PBS to ∼ 10^7^ particles mL^-1^ before measurements. The data of size distribution were analyzed with NTA software. The measurements were conducted at 25 °C.

### TEM

10 μL sEV sample (∼ 10^12^ particles mL^-1^) were directly absorbed on Formvar/carbon-coated copper grids for two minutes. After blotting residual samples with filter paper, 10 μL 1% phosphotungstic acid was dropped to stain sEV for 45 seconds followed by blotting the phosphotungstic acid with filter paper. After drying at room temperature, the grids containing sEV were observed on Tecnai G2 spirit Biotwin TEM (FEI) at 80 kV. For immunogold labeling of sEV sample derived from LNCaP, CD63 aptamer-conjugated gold nanoparticles were incubated with sEV samples for 30 min at 4 °C, and the samples were dropped in grids for TEM detection. 10 uL CD63 aptamer (1uM) was mixed with 100 uL gold nanoparticles (4 ∼ 10 nm, 2 mg/mL) at 4 °C for 12 h to prepare CD63 aptamer-conjugated gold nanoparticles, followed by blocking active sites with BSA solution (1% w/v).

### Immunoblot analysis of sEV samples

Isolated sEV samples and cells were treated with radio immunoprecipitation assay (RIPA) lysis buffer including protease inhibitors (Beyotime) with an ice bath for 30 min, followed by quantifying with a BCA assay. Protein lysates were separated by sodium dodecyl sulfate-polyacrylamide gel electrophoresis (SDS-PAGE), followed by transferring to polyvinylidene fluoride (PVDF) membrane. The transferred PVDF membranes were blocked using 5% non-fat dry milk in TBST buffer (TBS powder, Servicebio, 0.05% Tween-20) at room temperature for 1 h. Then blocked membranes were immunoblotted with a panel of primary antibodies including anti-Alix (Santa Cruz, sc-7964), anti-Tsg101 (Santa Cruz, sc-7964), anti-Calnexin (Abcam, ab133615), anti-CD63 (NOVUS, NBP2-4225B), anti-PTK7 (BBI, D199285-0100), anti-PSMA (Abcam, ab79542), anti-HER-2 (Abcam, ab134190), anti-EpCAM (BBI, D263391) overnight at 4 °C. Followed by incubating with corresponding HRP-conjugated secondary antibody for 1 h at 37 °C, the membranes were washed three times for 10 min at room temperature with TBST buffer. Finally, the western blot images were recorded on a gel image system (Tanon).

### Data analysis

The expression levels of target markers were defined by normalizing the target-associated number recorded to those of total number of sEV. The total number of sEV was measured by counting the number of sEV binding to the positive-charged sensor surfaces. The protein levels of total sEV and three small-EVs subtypes were normalized by subtracting the 2.5th percentile value and dividing by (97.5th percentile value–2.5th percentile value). These normalized data were directly used for LDA-based classification. Z-score were calculated by protein levels subtracted by mean and divided by row standard deviation. The significance of the difference between the sEV from cancer cell lines and normal cell line using individual protein marker was calculated using a two-tailed, heteroscedastic t-test (**Fig. 3c**). The significance of the difference between the sEV from five cell lines using individual protein marker was calculated using a two-tailed, heteroscedastic

### The artificial intelligence algorithm

The artificial intelligence algorithm developed here is based on Python 3.7. Because the sizing accuracy of iPM system is 10 nm, the data of SEVEN between 30 to 160 nm were re-calculated into thirteen subgroups at the interval of 10 nm. Theoretically, there are totally 2^12^ possibilities of subsets we can choose. Hill climbing algorithm was a random local search optimization algorithm for nonlinear objective functions, which was used to search for the locally optimal solution. Basically, hill climbing algorithm includes the following steps. 1) a group of subsets was randomly set as the initial point and its classification accuracy calculated by LDA was set as current best solution. 2) the next new point was defined by the greedy strategy of hill climbing algorithm. And the new classification accuracy was obtained. 3) the new classification accuracy was compared with the previous one. If the new classification accuracy was equal to or bigger than the previous one, the former point was abandoned and the latter was set as the current point, and vice versa. Then repeat step 2 and step 3 to quickly find the optimal solution.

## Author contribution

H.Y. and Y.Y. conceived the idea; C.Z. and Y.Y. performed the experiments; C.Z. and F.X. analyzed data; W.Z., Q.Z., J.W. and X.D. helped with instrumentation; H.H. and X.D. provided samples and helped with discussion; C.Z. and H.Y. wrote the paper.

## References

1. van Niel, G., D’Angelo, G. & Raposo, G. Shedding light on the cell biology of extracellular vesicles. Nat Rev Mol Cell Biol 19, 213–228 (2018).

2. LeBleu, V.S. & Kalluri, R. Exosomes as a multicomponent biomarker platform in cancer. Trends Cancer 6, 767–774 (2020).

3. Mathieu, M., Martin-Jaular, L., Lavieu, G. & Thery, C. Specificities of secretion and uptake of exosomes and other extracellular vesicles for cell-to-cell communication. Nat Cell Biol 21, 9–17 (2019).

4. Thery, C., Zitvogel, L. & Amigorena, S. Exosomes: composition, biogenesis and function. Nat Rev Immunol 2, 569–579 (2002).

5. Hoshino, A. et al. Tumour exosome integrins determine organotropic metastasis. Nature 527, 329–335 (2015).

6. Wortzel, I., Dror, S., Kenific, C.M. & Lyden, D. Exosome-mediated metastasis: communication from a distance. Dev Cell 49, 347–360 (2019).

7. Gyuris, A. et al. Physical and molecular landscapes of mouse glioma extracellular vesicles define heterogeneity. Cell Rep 27, 3972–3987 (2019).

8. Hoshino, A. et al. Extracellular vesicle and particle biomarkers define multiple human cancers. Cell 182, 1044–1061 e1018 (2020).

9. Kalluri, R. & LeBleu, V.S. The biology, function, and biomedical applications of exosomes. Science 367 (2020).

10. Giulietti, M. et al. Exploring small extracellular vesicles for precision medicine in prostate cancer. Front Oncol 8, 221 (2018).

11. Whiteside, T.L. Validation of plasma-derived small extracellular vesicles as cancer biomarkers. Nat Rev Clin Oncol 17, 719–720 (2020).

12. Liu, C. et al. lambda-DNA-and aptamer-mediated sorting and analysis of extracellular vesicles. J Am Chem Soc 141, 3817–3821 (2019).

13. Tian, F. et al. Protein analysis of extracellular vesicles to monitor and predict therapeutic response in metastatic breast cancer. Nat Commun 12, 2536 (2021).

14. Liu, C. et al. Low-cost thermophoretic profiling of extracellular-vesicle surface proteins for the early detection and classification of cancers. Nat Biomed Eng 3, 183–193 (2019).

15. Zhang, H. et al. Identification of distinct nanoparticles and subsets of extracellular vesicles by asymmetric flow field-flow fractionation. Nat Cell Biol 20, 332–343 (2018).

16. Bordanaba-Florit, G., Royo, F., Kruglik, S.G. & Falcon-Perez, J.M. Using single-vesicle technologies to unravel the heterogeneity of extracellular vesicles. Nat Protoc 16, 3163–3185 (2021).

17. Kowal, J. et al. Proteomic comparison defines novel markers to characterize heterogeneous populations of extracellular vesicle subtypes. Proc Natl Acad Sci U S A 113, E968–977 (2016).

18. Zheng, H. et al. Deconstruction of heterogeneity of size-dependent exosome subpopulations from human urine by profiling N-glycoproteomics and phosphoproteomics simultaneously. Anal Chem 92, 9239–9246 (2020).

19. Guan, S. et al. Size-dependent sub-proteome analysis of urinary exosomes. Anal Bioanal Chem 411, 4141–4149 (2019).

20. Wu, M. et al. Isolation of exosomes from whole blood by integrating acoustics and microfluidicss. Proc Natl Acad Sci U S A 117, 28525 (2020).

21. Zhang, H. & Lyden, D. Asymmetric-flow field-flow fractionation technology for exomere and small extracellular vesicle separation and characterization. Nature Protocols 14, 1027–1053 (2019).

22. Chen, Y. et al. Exosome detection via the ultrafast-isolation system: EXODUS. Nat Methods 18, 212–218 (2021).

23. Hendrix, A. The nature of blood(y) extracellular vesicles. Nat Rev Mol Cell Biol 22, 243 (2021).

24. Yang, Y. et al. Interferometric plasmonic imaging and detection of single exosomes. Proceedings of the National Academy of Sciences 115, 10275–10280 (2018).

25. Yang, Y., Zhai, C., Zeng, Q., Khan, A.L. & Yu, H. Multifunctional detection of extracellular vesicles with surface plasmon resonance microscopy. Analytical Chemistry 92, 4884–4890 (2020).

26. Thery, C., Amigorena, S., Raposo, G. & Clayton, A. Isolation and characterization of exosomes from cell culture supernatants and biological fluids. Curr Protoc Cell Biol Chapter 3, Unit 3 22 (2006).

27. Thery, C. et al. Minimal information for studies of extracellular vesicles 2018 (MISEV2018): a position statement of the International Society for Extracellular Vesicles and update of the MISEV2014 guidelines. J Extracell Vesicles 7, 1535750 (2018).

28. Zhou, S. et al. Accurate cancer diagnosis and stage monitoring enabled by comprehensive profiling of different types of exosomal biomarkers: surface proteins and miRNAs. Small 16, e2004492 (2020).

29. Xiao, X. et al. Intelligent Probabilistic System for Digital Tracing Cellular Origin of Individual Clinical Extracellular Vesicles. Anal Chem 93, 10343–10350 (2021).

