## Supplemental FigureS1-11 and Table S1 for "Profiling the size-dependent heterogeneity of membrane proteins in a mixed population of small extracellular vesicle for potential cancer diagnosis"

**This PDF file includes:**

Supplementary Text

Figs. S1 to S11

Tables S1

References (1 to 18)

Supplementary Text

**1. The surface marker selection**

CD63 is a universal exosomal biomarker, which is mostly used in exosomes immunological detection methods(*1*). Our western blotting results (**Figure 2b**, **Figure S5**) showed a positive expression among the all five samples, indicating a large number of exosomes in the samples. EpCAM is overexpressed in various epithelial carcinoma, and it is widely used in exosomal detection. sEVs derived from A549, MCF-7, HepG2 and LNCaP showed positive expression of EpCAM, which is consistent with previous studies(*2*). In our experiments, the normal liver cell showed negative expression of EpCAM(*3*). HER2 is one of the ErbB family members, which is related to the occurrence and development of breast cancer as well as is an effective prognostic indicator of breast cancer. Besides, HER2 is a tyrosine kinase receptor, which is a potent cancer-related protein in various cancers such as lung (A549 cell lines)(*4, 5*), and prostate cancer(*6, 7*). The expression of HER2 was also detected in exosomes derived from liver cells, for example HepG2 and L-02(*8*). We also observed the expression of HER2 in the sEVs from all the five cell lines in our experiment (**Figure S5**). PSMA is a prostate cancer marker, which is highly expressed in LNCaP cell line and LNCaP sEVs(*9*). But PSAM has also been discovered to be expressed in non-small cell lung cancer(*10*), and in extracellular vesicles of MCF-7(*11*)and HepG2(*12, 13*). In the five cell lines we used in this work, PSMA is positive in all samples. PTK7 is a key transmembrane receptor protein, which plays an essential role in regulating the Wnt signaling network(*14*). A549, MCF-7, LNCaP cells do not express PTK7(*15, 16*), which is consistent with our western blot results. The other two cell lines, HepG2 and L-02 expressed PTK7(*17*).

**2. Supplementary Figures**

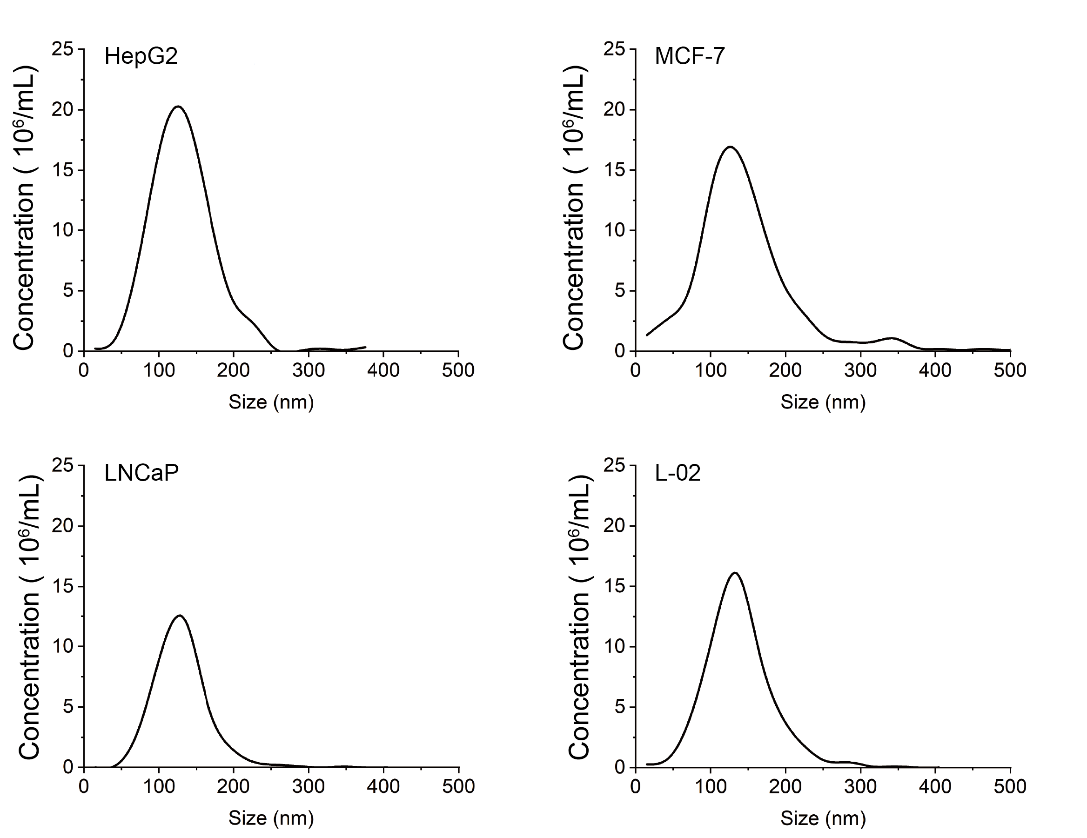

**Figure S1**. NTA measurement of HepG2, MCF-7, LNCaP and L-02 derived sEV samples

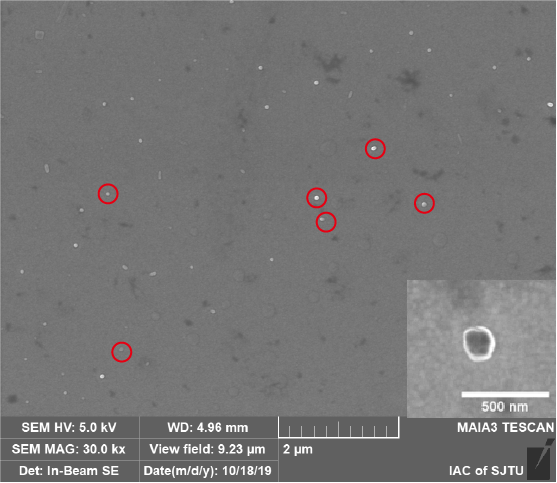

**Figure S2** TEM images of small EVs binding onto HER-2-modified chips.

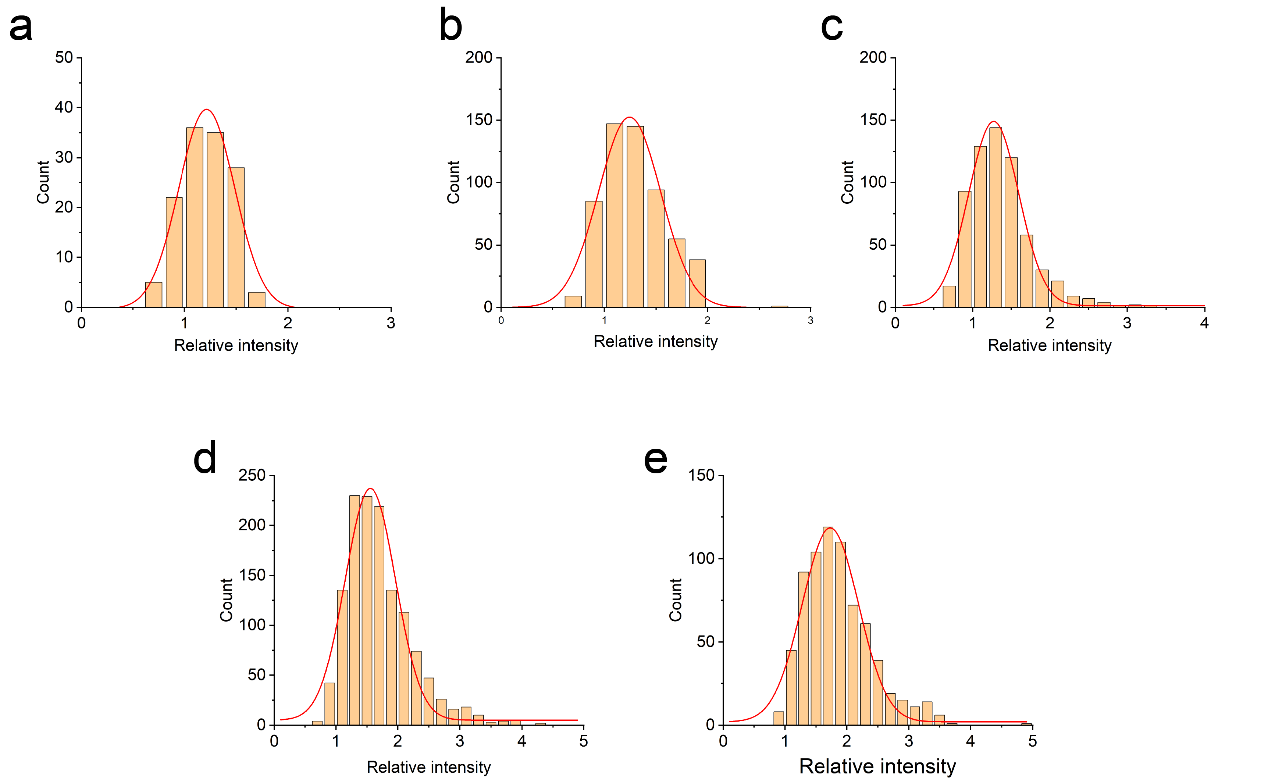

**Figure S3**. The iPM intensity change induced by single silica nanoparticles with diameters of (a) 30 nm (b), 50 nm, (c) 70 nm, (d) 100 nm and (e) 160 nm.

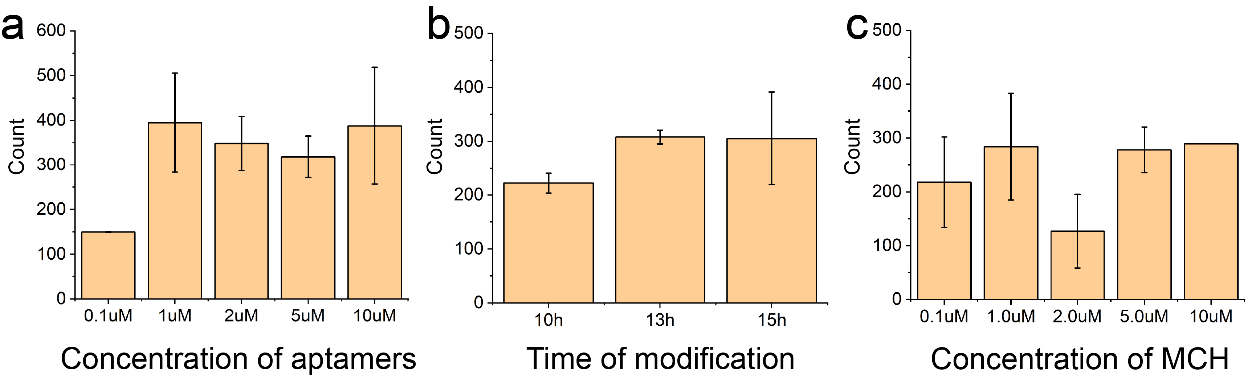

**Figure S4**. Optimization of surface modification. The recorded small EV (derived from MCF-7 cell line) number vs. (a) the concentration of CD63 aptamers, (b) the time of modification, and (c) the concentration of MCH.

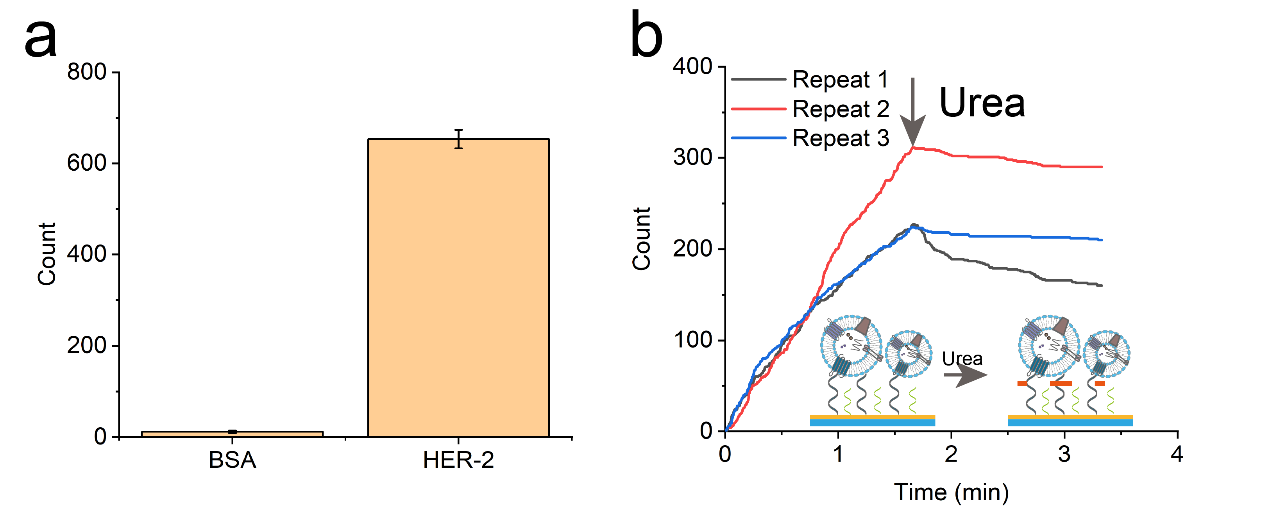

**Figure S5.** The specificity in small EV detection. (a) The number of small EVs recorded on BSA and HER-2 modified sensor surface, (b) the binding curves and competing tests for with urea.

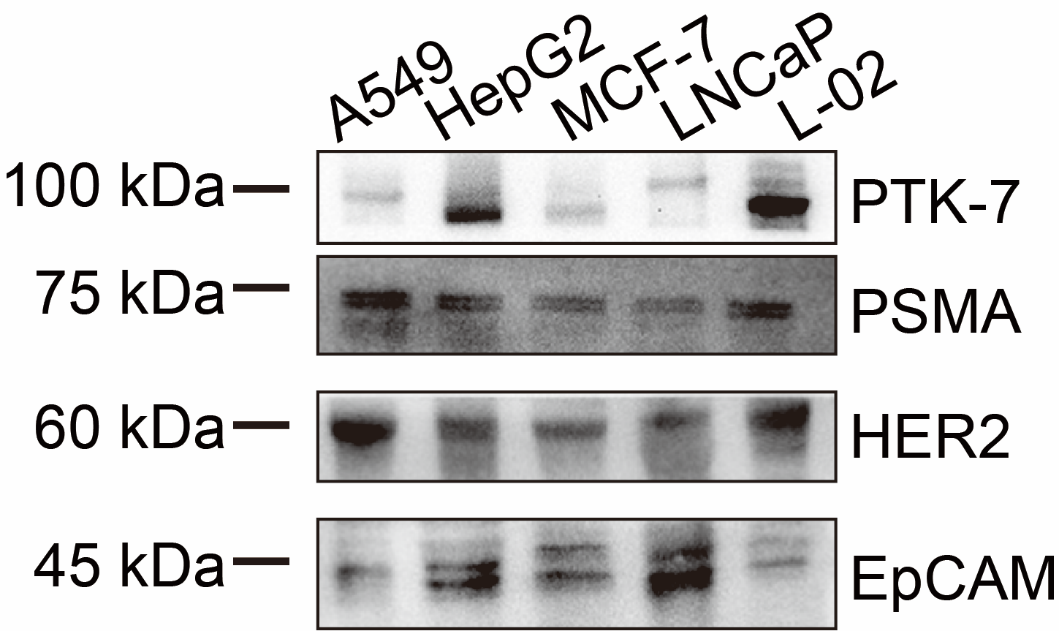

**Figure S6**. Western blot characterization of A549-, HepG2-, MCF-7-, LNCaP- and L-02- derived small EV samples.

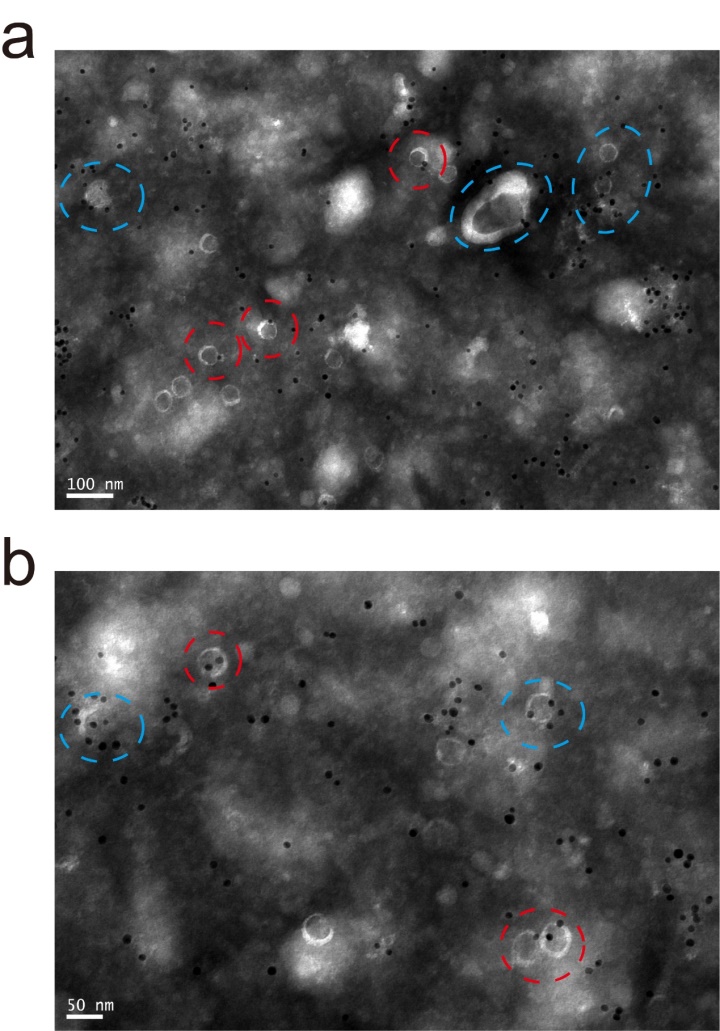

**Figure S7.** Immunoelectron microscopic image of the CD63-aptamer-coated gold nanoparticles on small EVs.

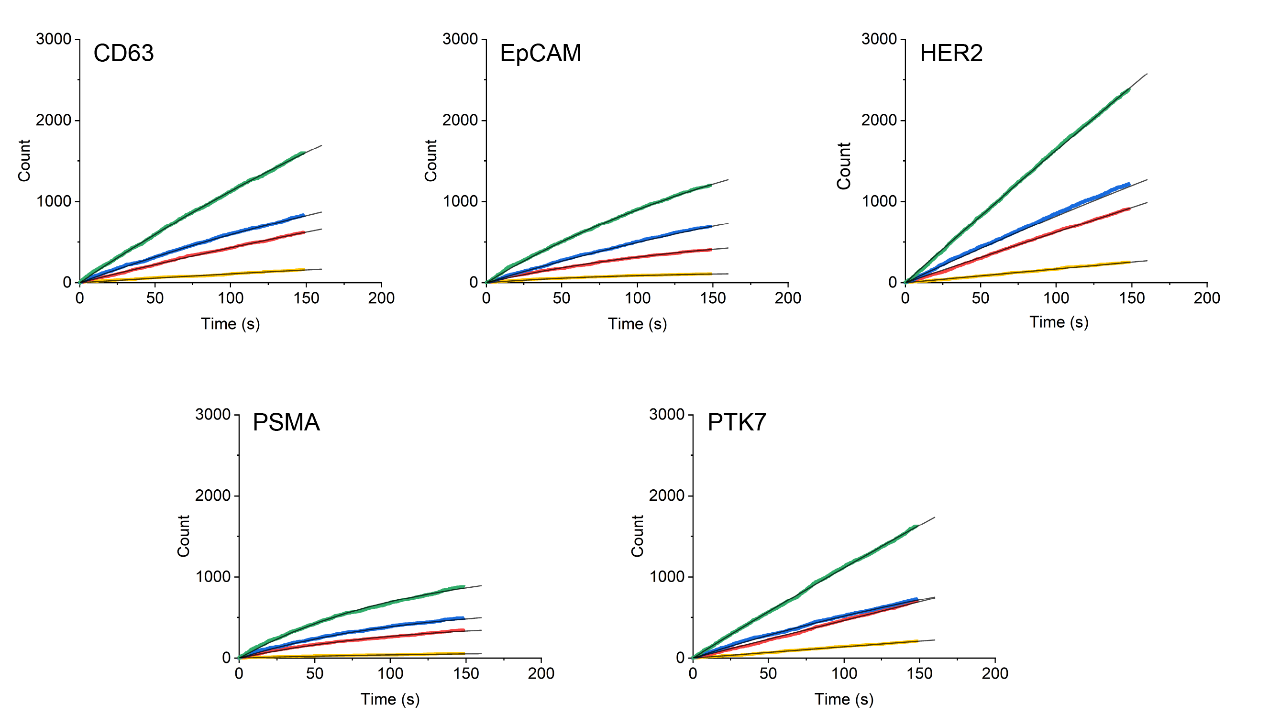

**Figure S8**. The binding kinetics of LNCaP-derived small EVs on aptamers modified sensor chips.

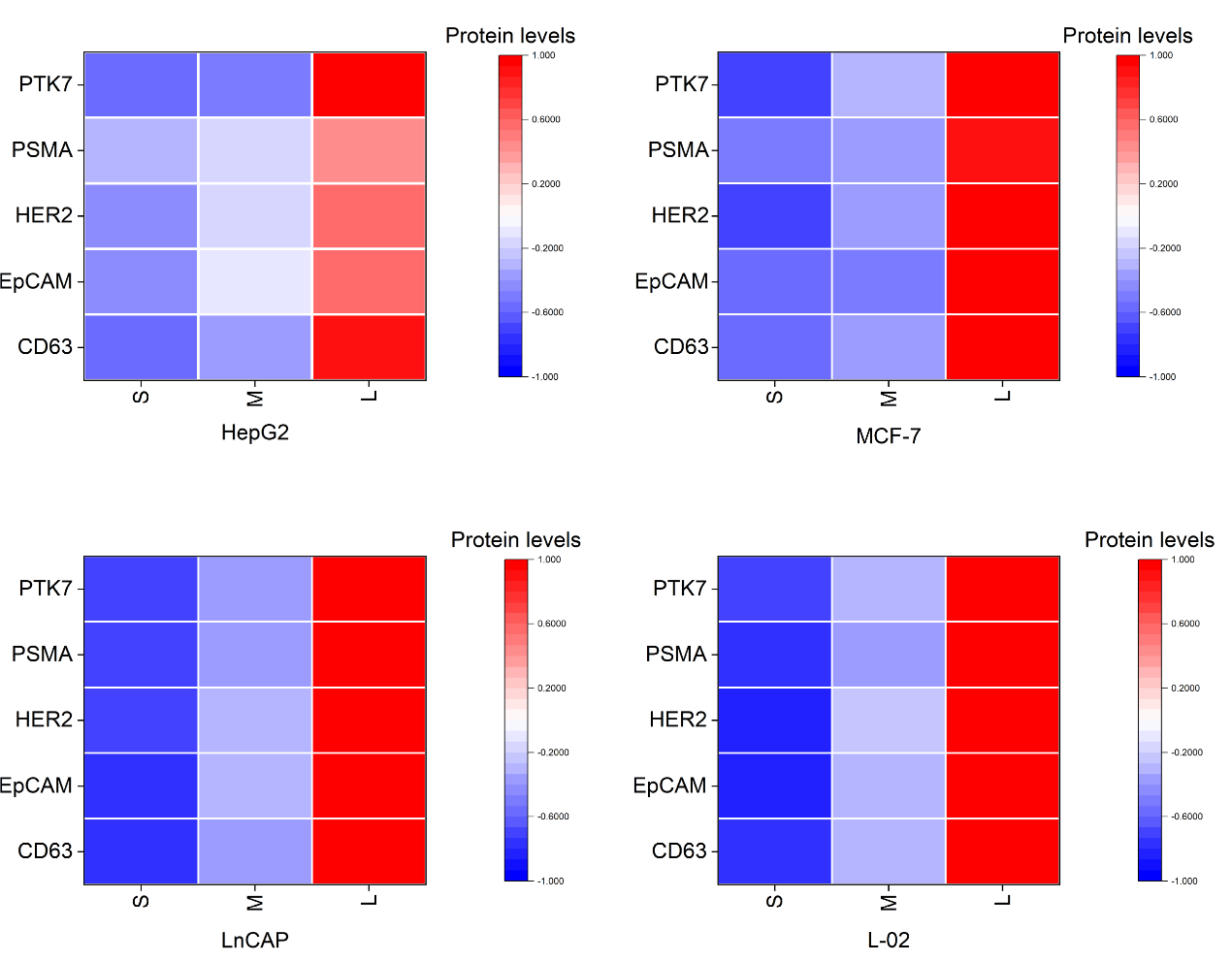

**Figure S9**. Heatmap illustration of the relative abundance of small EVs (HepG2, MCF-7, LNCaP, L-02) markers in sEV-S, sEV-M and sEV-L. Scale shown is protein levels subtracted by mean and divided by row standard deviation (that is, Δ (protein levels − mean)/s.d.).

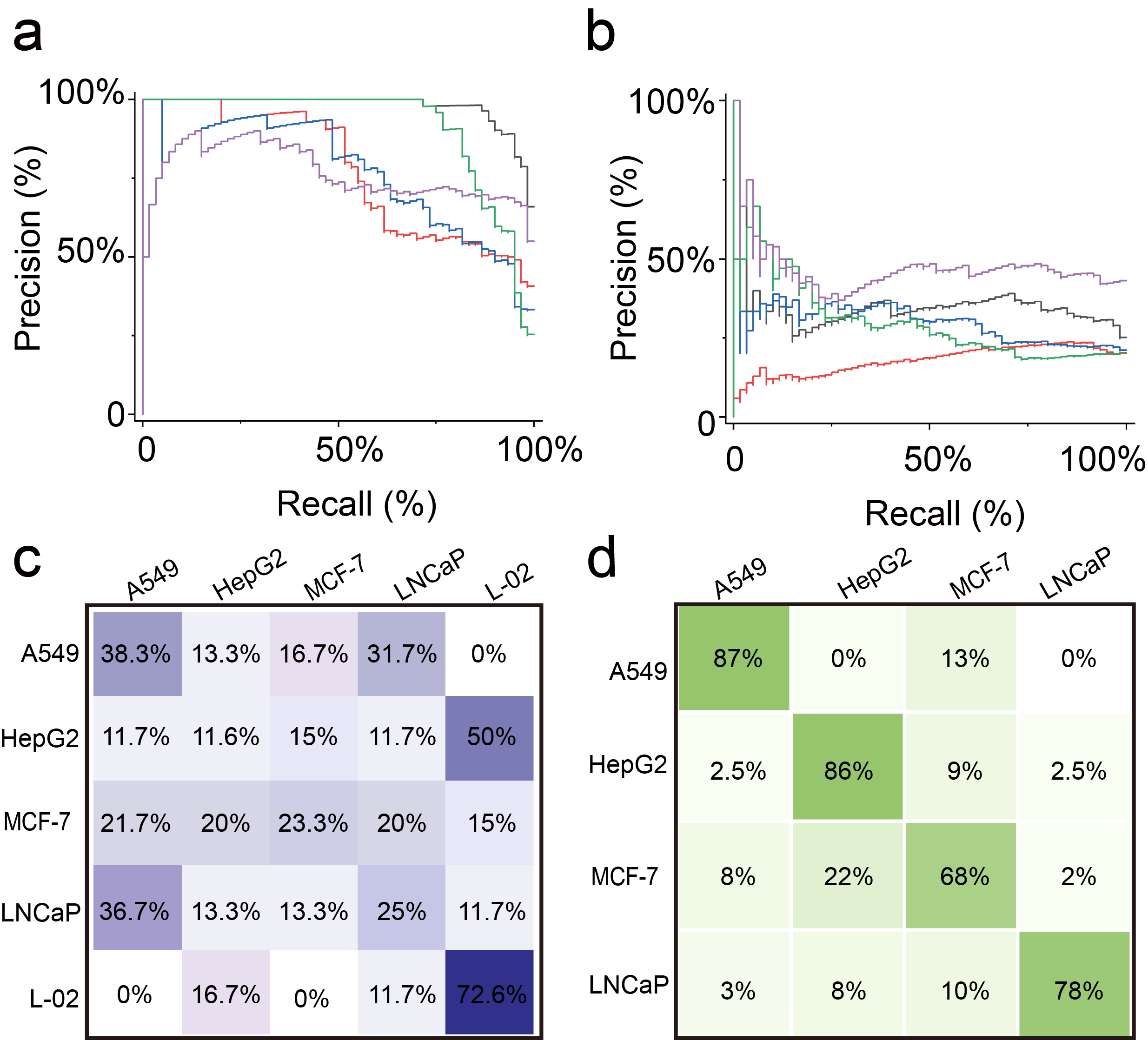

**Figure S10**. The cancer classification results. (a) Precision-Recall Curves (PRC) for multidimensional matrixes to differentiate five cancer cell lines with and (b) without subtype information. (c) Probability matrix summarizing the cancer classification results of multidimensional matrixes without size-dependent heterogeneous information. (d) Probability matrix of combining the data of HepG2 and L-02.

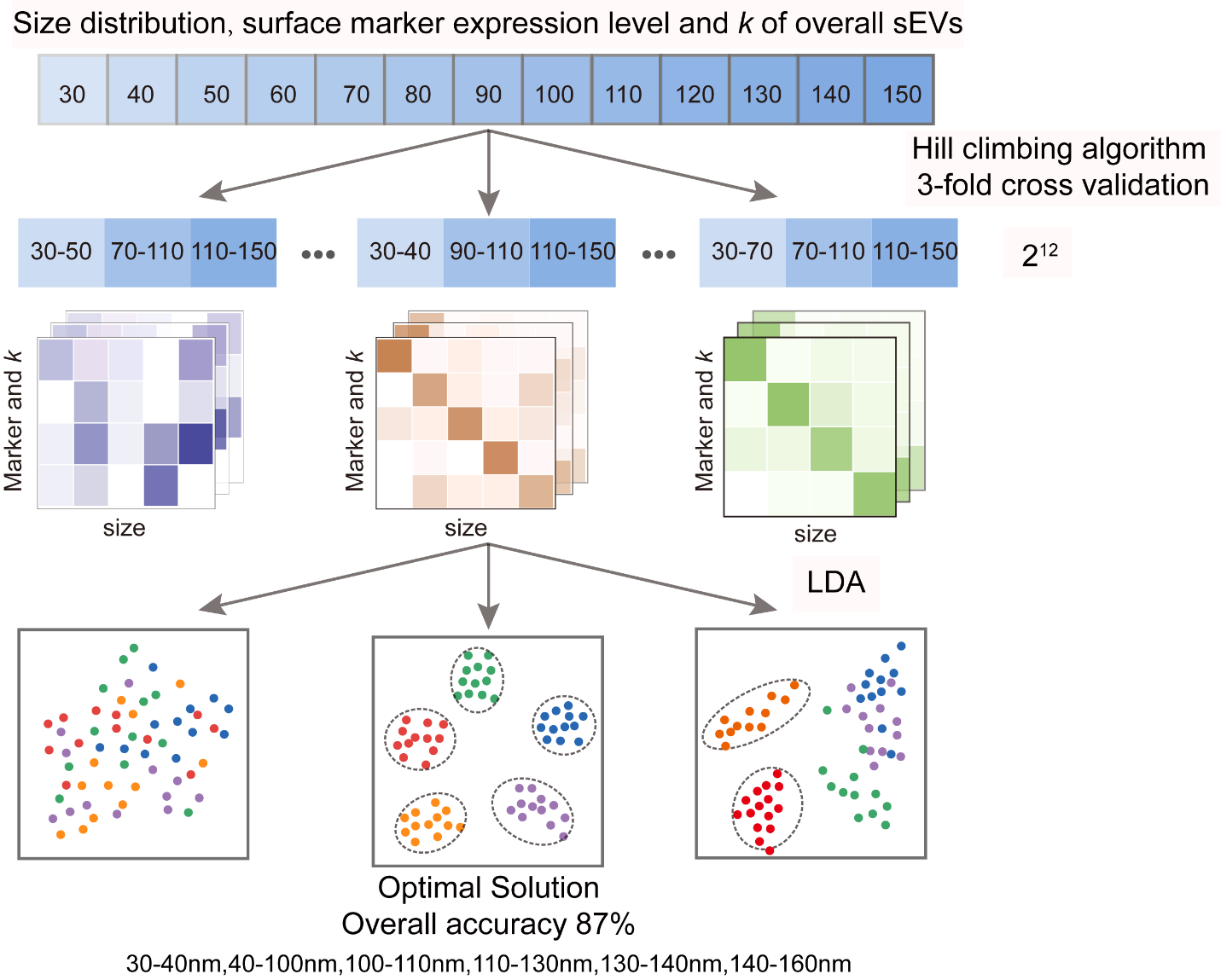

**Figure S11.** The schematic of Automatic Searching algorithm.

**Table S1. Summary of aptamers(*18*)**

| Aptamer | Sequence (5’ to 3’) |
| --- | --- |
| HER2 | 5'-GGG CCG TCG AAC ACG AGC ATG GTG CGT GGA CCT AGG ATG ACC TGA GTA CTG TCC-3' |
| EpCAM | 5'-CAC TAC AGA GGT TGC GTC TGT CCC ACG TTG TCA TGG GGG GTT GGC CTG-3' |
| PTK7 | 5'-ATC TAA CTG CTG CGC CGC CGG GAA AAT ACT GTA CGG TTA GA-3 |
| CD63 | 5'-CAC CCC ACC TCG CTC CCG TGA CAC TAA TGC TA-3' |
| PSMA | 5‘-GCG TTT TCG CTT TTG CGT TTT GGG TCA TCT GCT TAC GAT AGC AAT GCT-3‘ |

**References**

1.	N. Cheng *et al.*, Recent advances in biosensors for detecting cancer-derived exosomes. *Trends Biotechnol* **37**, 1236-1254 (2019).

2.	Y. Fan *et al.*, High-sensitive and multiplex biosensing assay of NSCLC-derived exosomes via different recognition sites based on SPRi array. *Biosens Bioelectron* **154**, 112066 (2020).

3.	M. Zhu *et al.*, HBx drives alpha fetoprotein expression to promote initiation of liver cancer stem cells through activating PI3K/AKT signal pathway. *Int J Cancer* **140**, 1346-1355 (2017).

4.	P. A. Bunn *et al.*, Expression of Her-2/neu in human lung cancer cell lines by immunohistochemistry and fluorescence in situ hybridization and its relationship to in vitro cytotoxicity by trastuzumab and chemotherapeutic agents. *Clinical Cancer Research* **7**, 13 (2001).

5.	T. Y. Rakovich *et al.*, Highly sensitive single domain antibody-quantum dot conjugates for detection of HER2 biomarker in lung and breast cancer cells. *ACS Nano* **8**, 14 (2014).

6.	M. Dahl *et al.*, Sarcosine induces increase in HER2/neu expression in androgen-dependent prostate cancer cells. *Mol Biol Rep* **38**, 4237-4243 (2011).

7.	R. Jahanban-Esfahlan *et al.*, The herbal medicine Melissa officinalis extract effects on gene expression of p53, Bcl-2, Her2, VEGF-A and hTERT in human lung, breast and prostate cancer cell lines. *Gene* **613**, 14-19 (2017).

8.	H. Di *et al.*, Nanozyme-assisted sensitive profiling of exosomal proteins for rapid cancer diagnosis. *Theranostics* **10**, 9303-9314 (2020).

9.	B. Li *et al.*, Facile fluorescent aptasensor using aggregation-induced emission luminogens for exosomal proteins profiling towards liquid biopsy. *Biosens Bioelectron* **168**, 112520 (2020).

10.	H. L. Wang *et al.*, Expression of prostate-specific membrane antigen in lung cancer cells and tumor neovasculature endothelial cells and its clinical significance. *PLoS One* **10**, e0125924 (2015).

11.	A. G. Wernicke *et al.*, Prostate-specific membrane antigen expression in tumor-associated vasculature of breast cancers. *APMIS* **122**, 482-489 (2014).

12.	S. S. Chang *et al.*, Five different anti-prostate-specific membrane antigen (PSMA) antibodies confirm PSMA expression in tumor-associated neovasculature. *Cancer Res* **59**, 7 (1999).

13.	Y. Yu *et al.*, Engineering of exosome-triggered enzyme-powered DNA motors for highly sensitive fluorescence detection of tumor-derived exosomes. *Biosens Bioelectron* **167**, 112482 (2020).

14.	J. Chen *et al.*, Organization of protein tyrosine kinase-7 on cell membranes characterized by aptamer probe-based STORM imaging. *Anal Chem* **93**, 936-945 (2021).

15.	D. Fan *et al.*, A polydopamine nanosphere based highly sensitive and selective aptamer cytosensor with enzyme amplification. *Chem Commun (Camb)* **52**, 406-409 (2016).

16.	Y. Fan, L. Li, M. Lu, H. Si, B. Tang, In situ fluorescent profiling of living cell membrane proteins at a single-molecule level. *Chem Commun (Camb)* **55**, 4043-4046 (2019).

17.	J. Jiang, Y. Yu, H. Zhang, C. Cai, Electrochemical aptasensor for exosomal proteins profiling based on DNA nanotetrahedron coupled with enzymatic signal amplification. *Anal Chim Acta* **1130**, 1-9 (2020).

18.	C. Liu *et al.*, Low-cost thermophoretic profiling of extracellular-vesicle surface proteins for the early detection and classification of cancers. *Nat Biomed Eng* **3**, 183-193 (2019).
